# Phospholipid membranes promote the early stage assembly of α-synuclein aggregates

**DOI:** 10.1101/295782

**Authors:** Zhengjian Lv, Mohtadin Hashemi, Siddhartha Banerjee, Karen Zagorski, Jean-Christophe Rochet, Yuri L. Lyubchenko

## Abstract

Development of Parkinson’s disease is associated with spontaneous self-assembly of α-synuclein (α-syn). Efforts aimed at understanding this process have produced little clarity and the mechanism remains elusive. We report a novel effect of phospholipid bilayers on the catalysis of α-syn aggregation from monomers. We directly visualized α-syn aggregation on supported lipid bilayers using time-lapse atomic force microscopy. We discovered that α-syn assemble in aggregates on bilayer surfaces even at the nanomolar concentration of monomers in solution. The efficiency of the aggregation process depends on the membrane composition, being highest for a negatively charged bilayer. Furthermore, assembled aggregates can dissociate from the surface, suggesting that on-surface aggregation can be a mechanism by which pathological aggregates are produced. Computational modeling revealed that interaction of α-syn with bilayer surface changes the protein conformation and its affinity to assemble into dimers, and these properties depend on the bilayer composition. A model of the membrane-mediated aggregation triggering the assembly of neurotoxic aggregates is proposed.

## Introduction

Parkinson’s disease (PD) is a devastating neurodegenerative disorder associated with the presence of cytosolic inclusions, named Lewy bodies (LBs), that contain amyloid-type fibrils typically localized to vertebrate presynaptic terminals (Goedert et al, 2013). These fibrils are assembled from α-synuclein (α-syn) protein, which is capable of spontaneous self-assembly into amyloid aggregates in solution (Luth et al, 2015). In addition to PD, α-syn is associated with the development of several neurodegenerative diseases, including LB Dementia and Alzheimer’s disease; however, the aggregation mechanism remains elusive (Ahmed et al, 2010; Friedrich et al, 2010; Ross & Poirier, 2004).

The self-assembly process of amyloid proteins, including α-syn, has led to the amyloid cascade hypothesis (ACH), which posits that the spontaneous assembly of amyloidogenic polypeptides is the key feature that defines the disease state (Hardy, 1992; Hardy & Higgins, 1992; Karran et al, 2011). With numerous data supporting this hypothesis, it remains the one on which *in vitro* and *in vivo* studies related to the molecular mechanisms of amyloid aggregation are based. However, there is a serious complication when translating current knowledge on amyloid aggregation *in vitro* to the aggregation process *in vivo* — namely, the concentrations of amyloidogenic polypeptide are dramatically different *in vivo* versus *in vitro*. The critical concentration for the spontaneous aggregation of α-syn *in vitro* is in the high micromolar range, in stark contrast to the nanomolar concentration range of α-syn in the cerebrospinal fluid (CSF) (Wang et al, 2012). We have discovered a novel aggregation pathway that essentially eliminates the concentration issue with ACH. We were able to observe spontaneous assembly of Aβ peptides and α-syn proteins at the nanomolar concentration range (Banerjee et al, 2017). This novel pathway is an on-surface aggregation mechanism that allows Aβ peptides of different sizes and α-syn proteins to assemble into aggregates at the nanomolar range. The process takes place at ambient conditions, physiological pH values, and with no agitation. Based on our finding, we hypothesized that similar on-surface aggregation mechanism was possible on membrane surfaces.

Interaction of α-syn aggregates with cellular membranes accompanied by changes in the membrane’s properties is considered a mechanism of PD development (Pfefferkorn et al, 2012). One of the current models posits that neurotoxic effects are associated with membrane permeability and cell death mediated by interactions with oligomeric α-syn (Dante et al, 2008; Green et al, 2004; Quist et al, 2005; Stockl et al, 2013). Interestingly, monomeric α-syn interacts with membranes as part of its normal function through binding to phospholipid molecules (Davidson et al, 1998; Diao et al, 2013), a property that is neglected in current models involving the assembly of toxic aggregates in bulk solution. Past reports suggest that the normal membrane binding function of α-syn is related to regulation of synaptic vesicle trafficking (Davidson et al, 1998; Diao et al, 2013; Venda et al, 2010). The protein has also been shown to undergo accelerated aggregation when incubated in the presence of phospholipid vesicles at high protein:lipid ratios (Galvagnion et al, 2015a; Lee et al, 2002; Ysselstein et al, 2015). It is proposed that α-syn aggregation at the membrane surface is stimulated both by the exposure of hydrophobic residues as the membrane-bound protein shifts from the long-helix form to the short-helix form (Bodner et al, 2009; Ysselstein et al, 2015), and by the increased probability of molecular interactions needed for α-syn self-assembly to occur on the two dimensional surface of the lipid bilayer than in solution (Abedini & Raleigh, 2009). The key role that membrane-induced α-syn aggregation plays in neurotoxicity (Ysselstein et al, 2015), potentially via a mechanism involving membrane permeabilization (Comellas et al, 2012; Lee et al, 2012; Reynolds et al, 2011; Ysselstein et al, 2017), is in line with our recent finding that surface interactions dramatically facilitates the aggregation process of amyloidogenic proteins, including α-syn. Together, these findings suggest that self-assembly at the surface of cellular membranes is the mechanism by which potentially neurotoxic oligomers are assembled at physiological concentration of the protein (Banerjee et al, 2017).

In the current study, this hypothesis is tested by direct visualization of α-syn aggregation on the surfaces of supported lipid bilayers (SLBs) using time-lapse atomic force microscopy (AFM). We demonstrate that SLBs catalyze the aggregation of α-syn at α-syn concentrations as low as 10 nM, which corresponds to the concentration range in the CSF (Wang et al, 2012). Aggregation kinetics are found to be dependent on SLB composition, being considerably higher for 1-palmitoyl-2-oleoyl-sn-glycero-3-phospho-L-serine (POPS) bilayer when compared to 1-palmitoyl-2-oleoyl-sn-glycero-3-phosphocholine (POPC). The assembled aggregates are not strongly bound to the surface and are capable of spontaneous dissociation into solution. Importantly, the self-assembly process does not damage the surface, as no defects were detected after the aggregate dissociation. Computational modeling demonstrates that α-syn monomers change conformation upon interaction with the bilayers and are dependent on bilayer composition. Conformations of α-syn after binding to POPS dramatically facilitate assembly of the dimer; a property in contrast with that on POPC and in line with experimental data. We propose a model for the membrane mediated amyloid aggregate assembly and the role of this process in beginning of the disease state.

## Results and Discussion

### Experimental AFM studies of α-syn aggregation on lipid bilayers

Lipid bilayers were assembled on the surface of freshly cleaved mica, using the approach described in the methods section, allowing for direct visualization of interactions between the protein and bilayer over many hours with AFM. Based on their prevalence in neuronal cellular membranes, three types of bilayers were used (Figure 1a): POPC, POPS, and an equimolar mixture of the two. The time-lapse AFM studies require that the bilayer is stable during the entire multi-hour AFM experiments. Additionally, the surface needs to be smooth, so protein aggregates can be detected as they are formed. We developed a procedure to assemble POPC and POPS bilayers, with no defects, over areas as large as 4 µm x 4 µm (Figure 1b and Appendix Figure S1). Each surface was tested for smoothness and stability prior to AFM studies of α-syn-membrane interactions. The stability and smoothness of the bilayer with the defect-free topography is illustrated in Appendix Figure S1. Only such smooth surfaces were used to conduct experiments.

**Figure 1:**
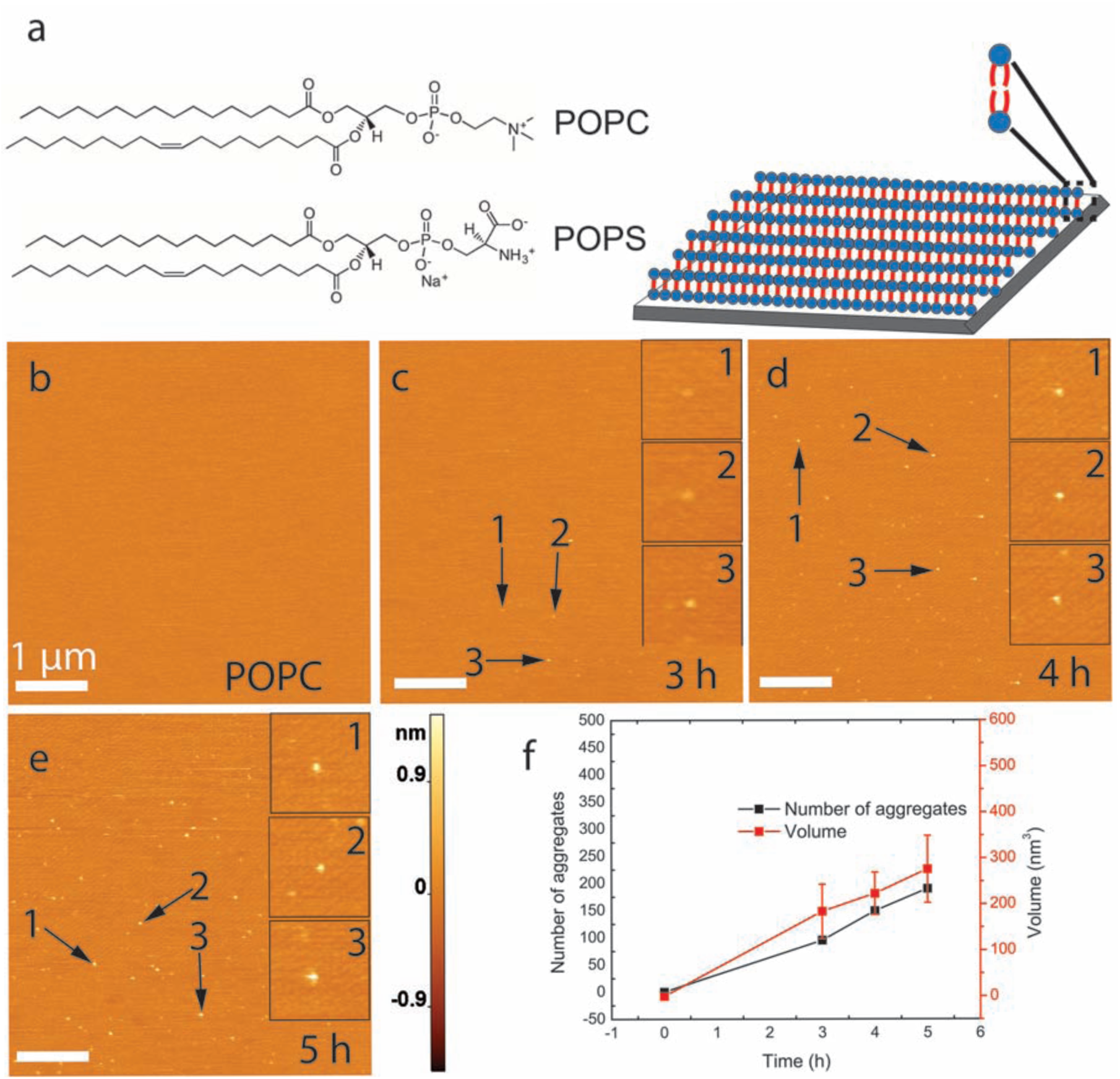
Time-lapse AFM images to characterize α-syn aggregation on supported POPC lipid bilayer (SLB). **(a)** POPC and POPS were used in the present study, and their chemical structures are shown (left). Schematic of a supported lipid bilayer (SLB) on freshly cleaved mica (right). The schematic is only for displaying the model for supported lipid bilayer, it does not indicate any phase of the bilayer. **(b)** Image of the POPC SLB surface immediately after buffer exchange with 10 nM α-syn solution. **(c)-(e)**, Images of SLB surface taken at time-points after adding the protein. Insets show zoomed images of three representative aggregates. **(f)** Graph showing the evolution of aggregate quantity and mean volume with respect to time. The data are shown as the mean ± SD. The scale bar in panels **(b)-(e)** is 1 μm, and the Z-scale is shown to the right of panel **(e)**.

After exchanging the buffer with a solution of α-syn, the bilayer was observed by time-lapse AFM imaging. Aggregation of α-syn on supported POPC bilayers was investigated over a period of 5 hours and the data is shown in Figure 1. Figure 1b shows the POPC surface immediately after exchange of buffer with 10 nM α-syn solution. Aggregates appear after 3 h of incubation, as indicated with arrows in Figure 1c. More aggregates appear over the next two hours of observation, and their sizes increase as well (Figures 1d-e). The number of aggregates and the aggregate sizes (volumes) were measured and are plotted in Figure 1f. The data shows that both parameters gradually increase over time. Similar time-lapse aggregation experiments were performed on APS-functionalized mica to compare the aggregation with aggregation on POPC SLB surface. The results are assembled as Appendix Figure S2. Qualitatively, aggregation on APS mica exhibits a similar trend to the POPC bilayer, with aggregates appearing after 3 h incubation (Appendix Figure S2e). However, the aggregation is less efficient as measured by the number and sizes of aggregates (Appendix Figures S2h and Figure EV1). Note that very few aggregates were found in a 10 nM solution of α-syn incubated under the same conditions in the absence of a phospholipid bilayer or mica surface.

To test the effect of the bilayer composition on α-syn aggregation, experiments were done with POPS bilayers. POPS shares hydrocarbon chains with POPC but has a serine head group that, at physiological pH, renders the surface negatively charged, unlike POPC which has a net neutral charge. AFM images in Figure 2 demonstrate that on POPS, aggregates appear after 1 h of incubation (Figure 2a), with the surface densely covered with α-syn aggregates after 5 h incubation (Figure 2b-d). Quantitative analysis demonstrated that the number of aggregates and their size increase over time (Figure 2e). The volume distribution histograms at each time point are shown in Figure EV1 (for more data see table 1). Compared with POPC, aggregation on POPS bilayers was accelerated, with the first detectable aggregates appearing after 1 h (POPS) vs 3 h (POPC). In addition, at the end of the experiment the number of aggregates was more than double for POPS (404) vs. POPC (190), and aggregates formed in the presence of POPS had a larger mean volume (417 nm^3^ for POPS vs 276 nm^3^ for POPC) (Figure EV1 and Table 1).

**Table 1:** Summary of results from three independent experiments. Volume and number of aggregates on POPC, POPS and POPC/POPS surfaces at different time intervals are shown.

|  |  | POPC |  | POPS |  | POPC/POPS |  |
| --- | --- | --- | --- | --- | --- | --- | --- |
|  |  | Volume<br>(nm<sup>3</sup>) | Aggregate<br>Number | Volume<br>(nm<sup>3</sup>) | Aggregate<br>Number | Volume<br>(nm<sup>3</sup>) | Aggregate<br>Number |
| Experiment 1 | 1 h | -- | -- | 143±41 | 129 | -- | -- |
|  | 2 h | -- | -- | 182±40 | 130 | 137±37 | 82 |
|  | 3 h | 184±59 | 95 | 255±94 | 141 | 187±59 | 176 |

**Table 1.**
|  |  |  |  |  |  |  |  |
| --- | --- | --- | --- | --- | --- | --- | --- |
|  | 4 h<br>5 h | 223±46<br>276±73 | 149<br>190 | 387±140<br>417±161 | 324<br>404 | 248±87<br>312±116 | 291<br>372 |
| Experiment 2 | 1 h | -- | -- | 123±48 | 116 | -- | -- |
|  | 2 h | -- | -- | 169±42 | 120 | 112±48 | 71 |
|  | 3 h | 159±53 | 86 | 227±91 | 272 | 176±73 | 166 |
|  | 4 h | 198±45 | 132 | 331±126 | 307 | 209±96 | 272 |
|  | 5 h | 228±85 | 172 | 363±165 | 383 | 289±136 | 351 |
| Experiment 3 | 1 h | -- | -- | 143±53 | 125 | -- | -- |
|  | 2 h | -- | -- | 212±47 | 140 | 157±46 | 91 |
|  | 3 h | 179±82 | 106 | 284±95 | 153 | 200±67 | 187 |
|  | 4 h | 240±41 | 161 | 414±142 | 343 | 277±83 | 283 |
|  | 5 h | 319±96 | 206 | 460±163 | 425 | 347±113 | 356 |

**Figure 2.**
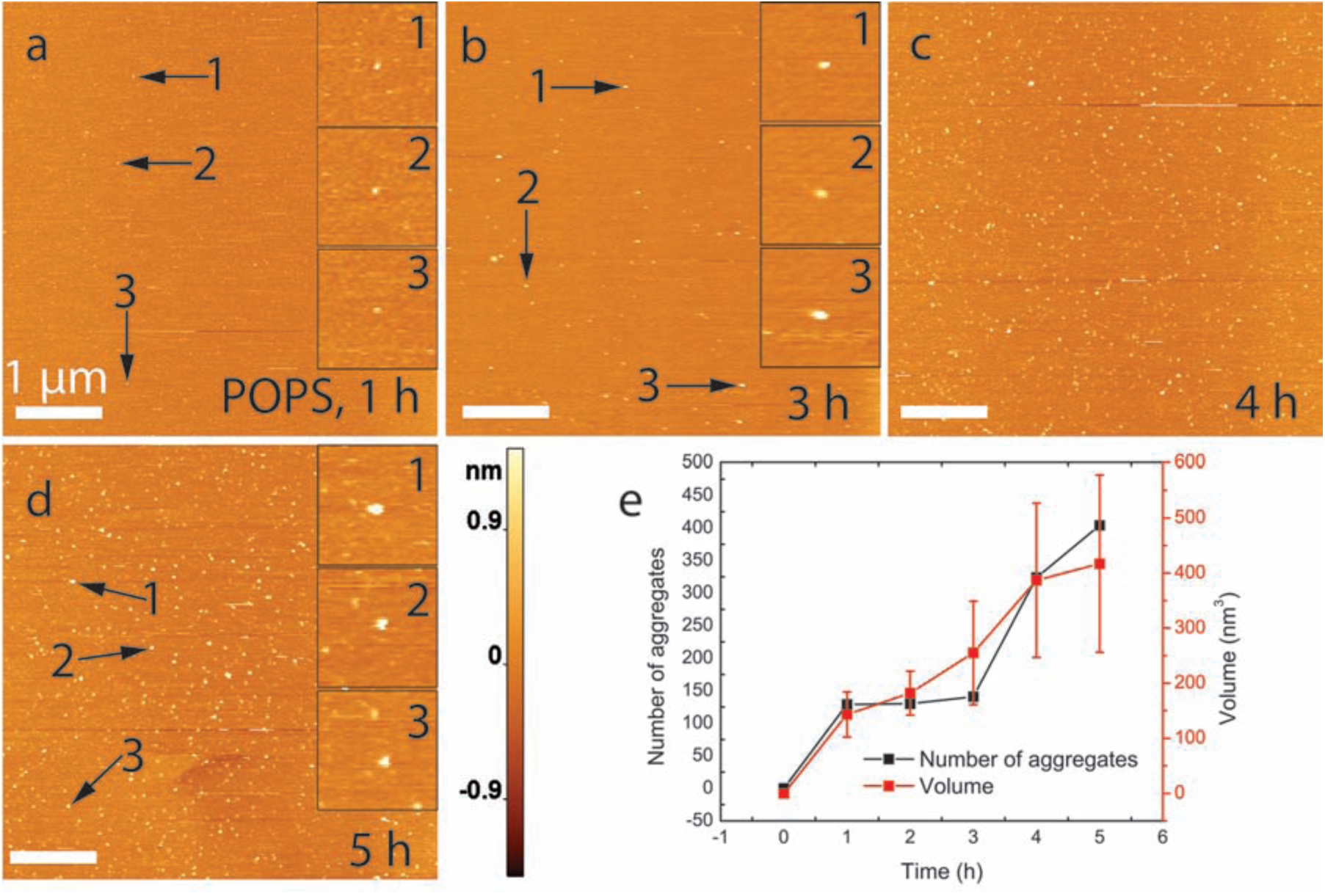
α-Syn shows enhanced aggregation kinetics on a supported POPS bilayer. **(a)-(d)** AFM images acquired at time-points after buffer exchange with 10 nM protein solution. Insets show zoomed images of three representative aggregates for the selected time-points. **(e)** Graph showing the time-dependent evolution of aggregate quantity and mean volume. The data are shown as the mean ± SD. The scale bar in panels **(a)-(d)** is 1 μm, and the Z-scale is shown to the right of panel **(d)**.

The aggregation of α-syn on a bilayer consisting of an equimolar mixture of POPC and POPS was the third bilayer composition tested. AFM data from the time-lapse experiments are shown in Figure 3. Aggregates appear after two hours (panel a), and their number increases over time (panels a-d). Quantitative analysis (Figure 3e) shows that the number and sizes of the aggregates grow monotonically with time. Although the number of aggregates on POPC/POPS is close to that on POPS (372 vs 404), the average size of aggregates was much smaller (see panel ‘Experiment 1’ in Table 1; mean volume of 312 nm^3^ on POPC/POPS vs 417 nm^3^ on POPS). Furthermore, the first appearance of aggregates on the equimolar mixture SLB is 1 h later than POPS SLB. The volume distributions at each time point for the POPC/POPS SLB is shown in Figure EV1 and Table 1. A direct comparison of aggregates on the different bilayer surfaces shows that the general aggregation propensity follows this order: POPS > POPC/POPS > POPC (Table 1).

**Figure 3.**
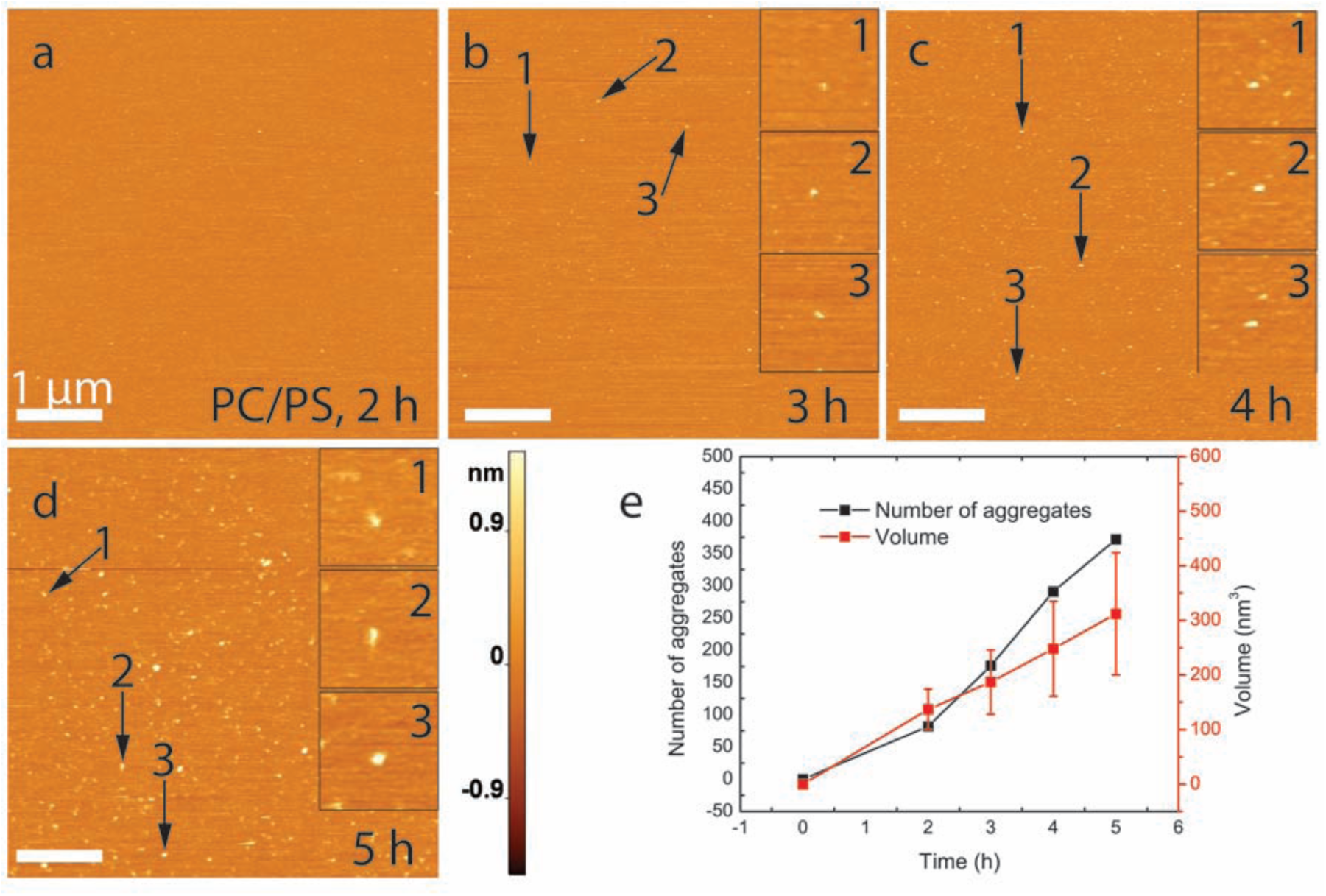
α-syn aggregation on a POPC/POPS SLB. **(a)** AFM image of initial bilayer acquired immediately after the exchange. **(b)-(d)** correspond to images taken with 1 h time intervals. Zoomed images of three representative aggregates are shown on the right side of images. **(e)** Line plot showing the time-dependent evolution of aggregate quantity and mean volume. The data are shown as the mean ± SD. The scale bar in panels **(a)-(d)** is 1 μm, and the Z-scale is shown to the right of panel **(d)**.

The data from time-resolved experiments allows us to follow the dynamics of individual aggregates. Images in Figure 4 are scans of the same area taken during a 30-minute interval. The aggregates are highlighted with arrows of different colors to indicate different types of aggregate dynamics. New aggregates appearing in panel B are highlighted with green arrows. The aggregates that do not change between panels A and B are marked with transparent black arrows. One aggregate on panel A dramatically increases in size in panel B and is highlighted with a yellow arrow. Interestingly, several aggregates highlighted in blue in panel A are missing in panel B, suggesting that these aggregates spontaneously dissociate from the surface during the 30-minute interval. Importantly, the surface remains smooth, indicating that no damage occurs to the bilayer surface following dissociation of the aggregates. Thus, aggregates assembled on the surface can dissociate back into solution, suggesting that aggregates should appear in the bulk solution above the bilayer. This assumption was tested by direct measurement of the time-dependent accumulation of α-syn aggregates in solution above the bilayer surface. To achieve this, aliquots were taken from the bulk solution (above bilayers), deposited on mica, imaged with AFM, and the aggregates analyzed. The results obtained for the sample taken in the presence and absence of SLBs are shown in Figure 5. The presence of the bilayer produces significantly more aggregates (solid black bars) compared to the control (dashed bars), supporting the conclusion about dissociating aggregates assembled on the lipid bilayer. Note that similar effect were observed in our recent paper (Banerjee et al, 2017), in which the assembly of α-syn aggregates on mica surface was studied.

**Figure 4.**
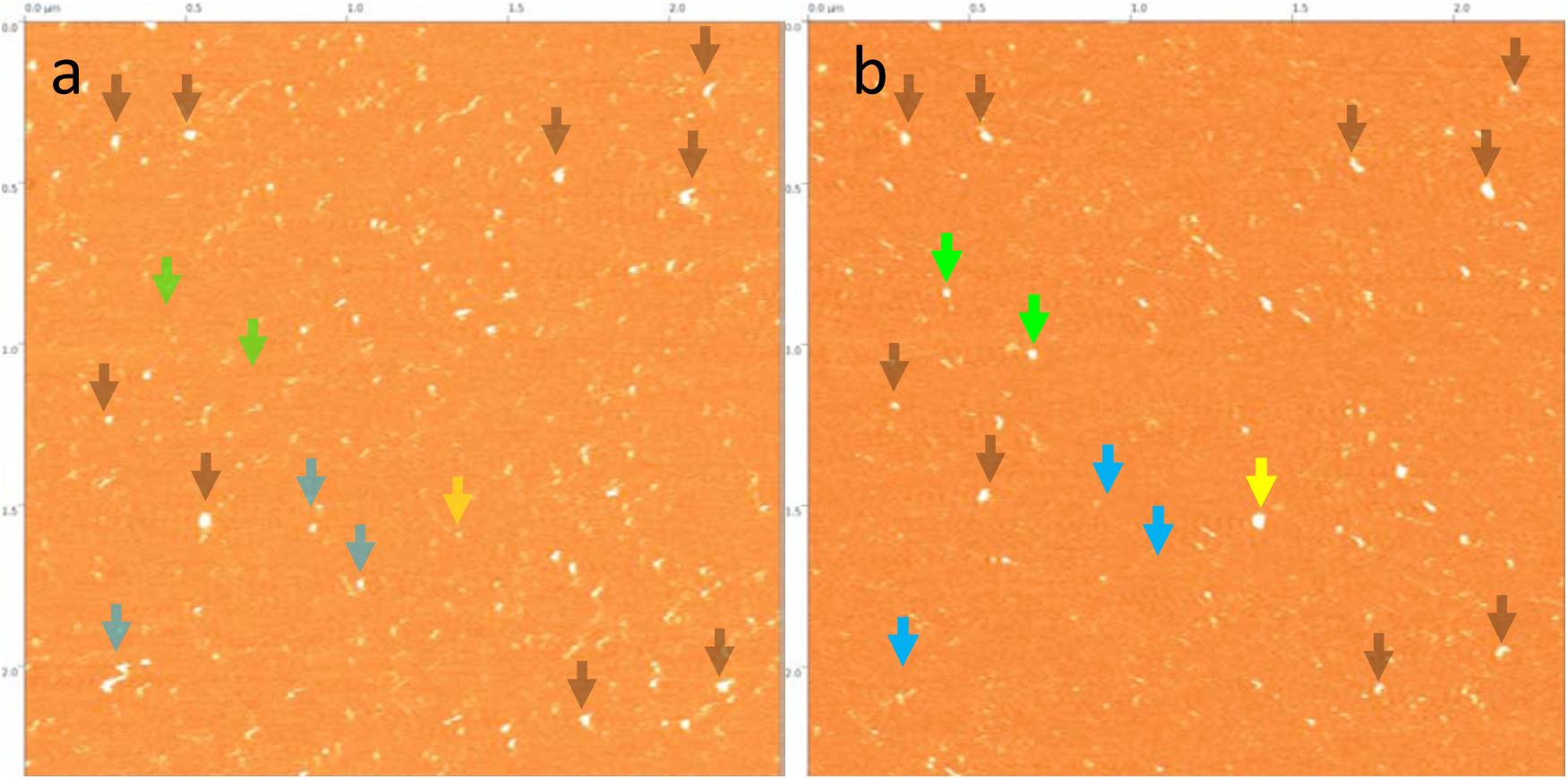
Dynamics of α-syn aggregates on POPS SLB captured after 6 h **(a)** and 6.5 h **(b)**. The aggregates highlighted with black arrows are features that did not change between frames and demonstrate the absence of drift. Blue arrows in panel **(a)** correspond to aggregates that have dissociated in panel **(b)**. New aggregates that appeared in **(b)** are highlighted with green. A growing aggregate is highlighted in yellow.

**Figure 5.**
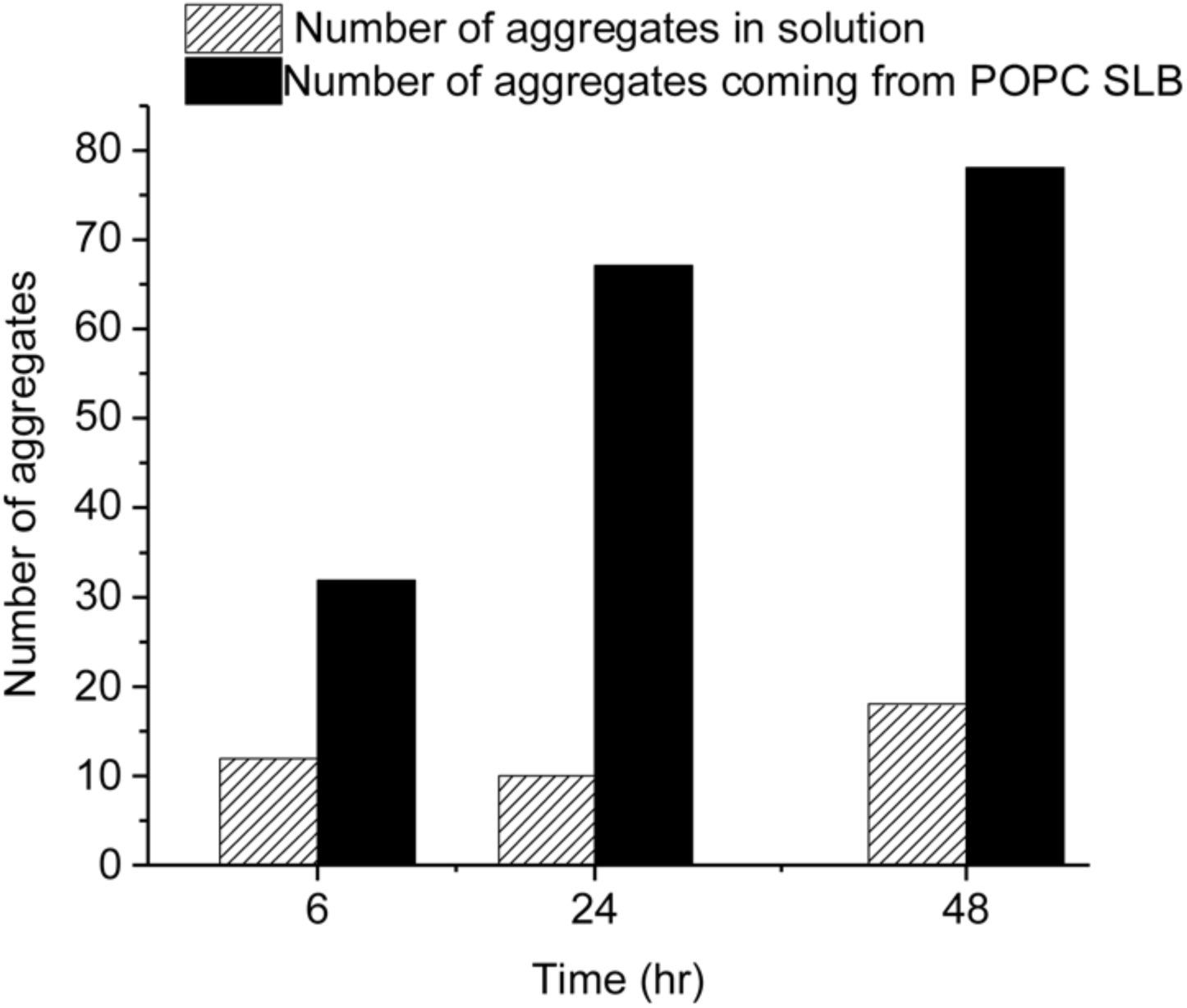
Accumulation of α-syn aggregates in solution from POPC SLB. A 10 nM α-syn solution was incubated in the presence of a POPC SLB. 5 µl of the solution was taken out at different time-points (6 h, 24 h, 48 h) and analyzed by AFM. Solid black bars show the number of aggregates, which appeared in the bulk from the POPC SLB, at different times. In a parallel experiment, a 10 nM α-syn solution was incubated in a test tube, and an aliquot of 5 µl was taken out at similar time-points and analyzed by AFM to check aggregation in the absence of a POPC SLB (striped bars). Aggregates were counted in 2 µm × 2 µm AFM images.

### Computational modeling of interaction of α-syn with lipid bilayers

To obtain insight into the underlying molecular mechanism of α-syn aggregation on the bilayer surface, we used molecular dynamics (MD) simulations of interaction of α-syn with the POPC and POPS bilayers. Briefly, a monomer of α-syn was placed 6 nm above the center of a 13 nm x13 nm bilayer patch (512 lipids), and interactions with the bilayer were then simulated. A few selected snapshots illustrating the dynamics of the interaction of α-syn with the POPC bilayer are shown in Figure 6a. The set of data for the interaction with POPC is assembled as Movie EV1. According to Figure 6a, α-syn initially binds to the POPC membrane through its N-terminal segment (frame (ii)). Over time, the length of the segment of the protein in contact with the POPC surface increases, so that the NAC segment approaches the surface as well (frames (iii-iv)). Graphically this change in binding is illustrated by the kymograph shown in Appendix Figure S3a. In fact, α-syn undergoes multiple association-dissociation events, as evidenced by the fluctuations of the number of contacts over time (Appendix Figure S3c). Eventually (after ~1.5 µs, seen as a jump in the graph), the protein strongly interacts with the bilayer and stays bound to the surface for the remainder of the simulation. Throughout the simulation the end-to-end distance and the radius of gyration of the α-syn molecule experience minor fluctuations (Appendix Figure S3d and e).

**Figure 6.**
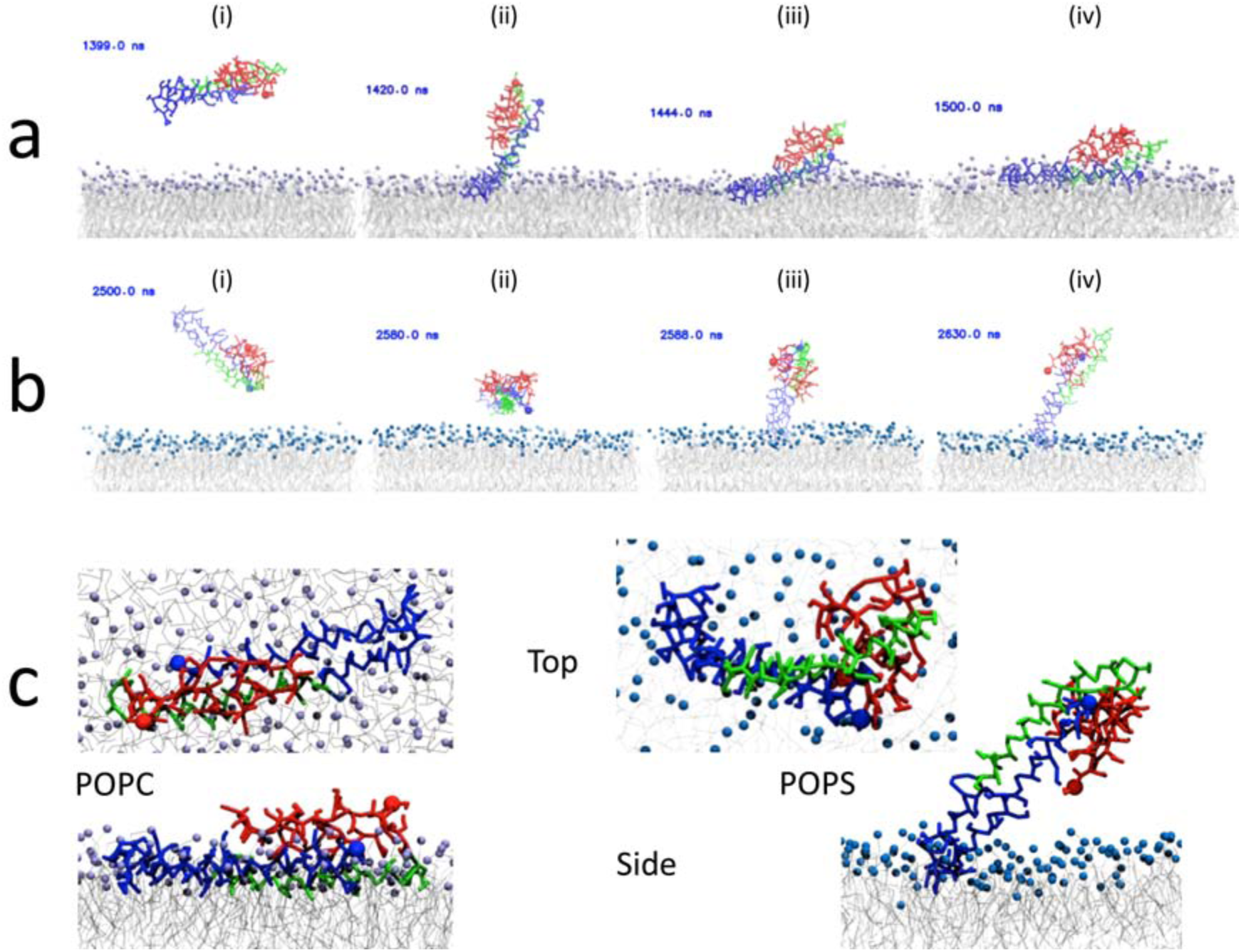
Molecular dynamics (MD) simulations of α-syn interaction with lipid bilayers reveal distinct conformations. The results show stable binding of α-syn to POPC **(a)** and POPS **(b)** bilayers; time-resolved stability is presented in Figure S6. **(c)** Top and side view snapshots of the last frame, at 4 μs, of the MD simulations for POPC and POPS, left and right respectively. The α-syn N-terminal segment is colored blue, the NAC region is in green, and the C-terminal segment is in red. N- and C-terminal residues are highlighted with a sphere. Lipid tails are in grey while the POPC and POPS head groups are in purple and blue, respectively.

A similar analysis was performed for the α-syn interaction with a POPS bilayer. A few selected frames are shown in Figure 6b, and the full set of data for the interaction is assembled as Movie EV2. Similar to the data obtained for the POPC bilayer, the N-terminal segment of α-syn binds to the membrane surface, but unlike POPC, the interaction with POPS is limited to a short central region (G36-K58) of the protein, graphically illustrated by Appendix Figure S3b. As a result, the protein remains extended out of the plane of the POPS surface.

We then modeled the interaction of membrane-bound α-syn with a second free α-syn molecule; the results are shown in Figure 7. Frame (i) in Figure 7a shows the second protein floating around the bound α-syn on POPC, but later (frame ii) it moves away from the bound protein and binds to the other side of the bilayer, gradually increasing the number of segments interacting with the bilayer as shown in frames (iii) to (iv). An animation of the dynamics is assembled as Movie EV3. Simulations with POPS bilayer produce entirely different results. According to Figure 7b, a free protein shown in frame (i) very rapidly binds to membrane-bound extended α-syn, and the dimer is formed rapidly after only 15 ns (frame (ii)) via interactions involving the two protein molecules’ NAC segments and via NAC-C-terminal interactions. The proteins in frame (iii) re-arrange the orientation to a parallel one with an extended NAC-NAC interaction interface (frames (iv)-(v)), and the dimer remains stable for the remainder of the simulation. The COM distance plot, Figure 7c, further corroborates these observations, with the distance on POPS quickly stabilizing to approximately 2.5 nm, while on POPC the distance is large fluctuating around 7 nm which is equal to the thickness of the bilayer plus contributions from the position on the bilayer surface (XY diffusion). The dynamic process is presented in Movie EV4. Furthermore, geometric analysis of the proteins, Appendix Figure S4, demonstrates that interactions within the dimer (on POPS) primarily occur between the NAC and C-terminal segments of the membrane-bound protein and the second α-syn molecule.

**Figure 7.**
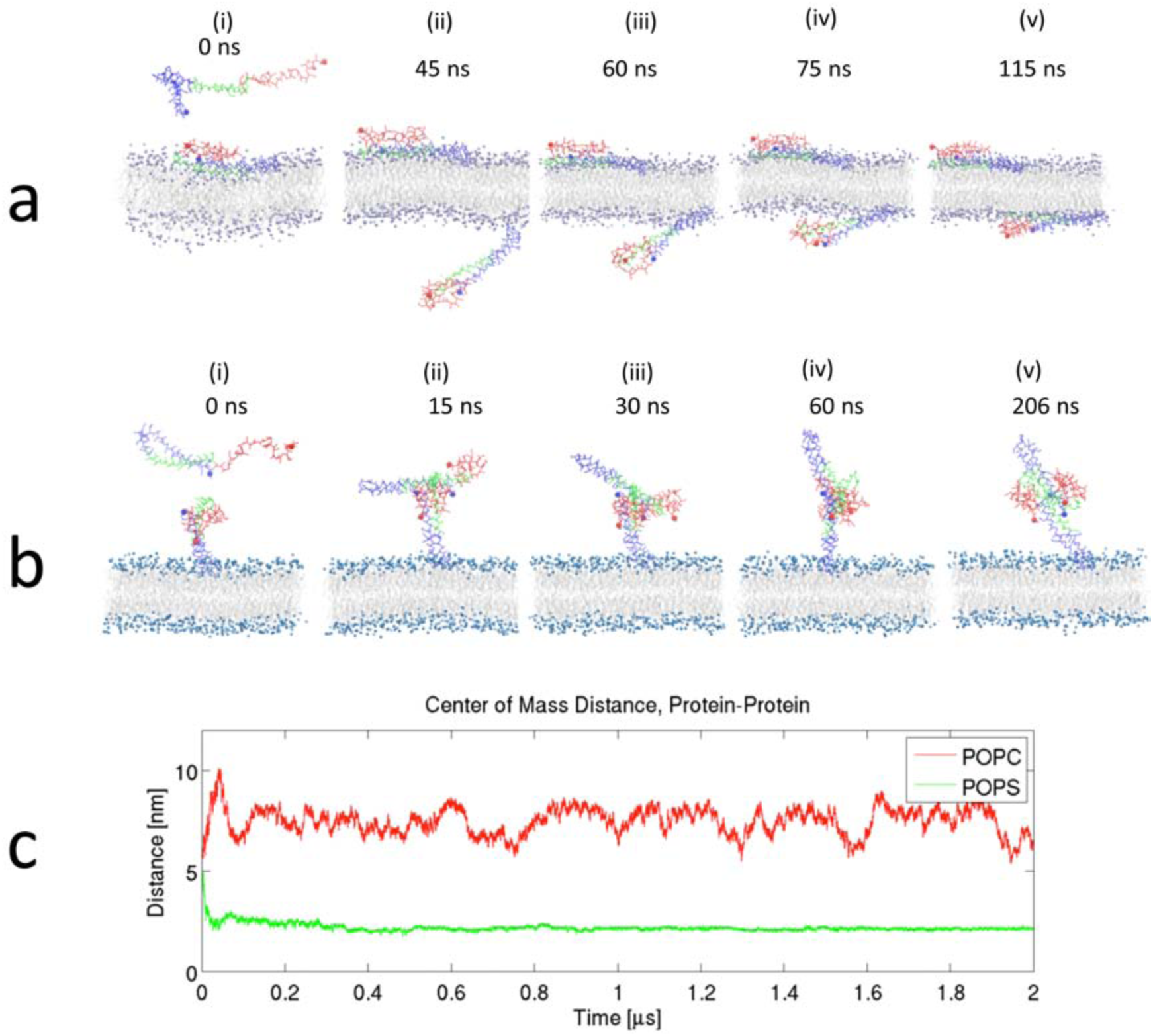
Results of MD simulations showing interaction between a free and a membrane-bound α-syn. **(a)** Binding of a free α-syn to the POPC membrane; the free α-syn traverses through the periodic boundary (PB) to the inner leaflet and stably binds; mode of interaction is similar to the initial α-syn interaction in Figure 4a. **(b)** On POPS the free α-syn rapidly binds membrane-bound α-syn through NAC-NAC and NAC-C-terminal interactions. **(c)** Center of mass (CoM) distance between the two α-syn molecules for the POPC and POPS systems. For POPC, after the transition through the PB, the fluctuations in CoM distance reflect the diffusion of the proteins in the XY-plane. The α-syn N-terminal segment is colored blue, the NAC region is in green, and the C-terminal segment is in red. N- and C-terminal residues are highlighted with a sphere. Lipid tails are in grey, while the POPC and POPS head groups are in purple and blue, respectively.

Overall, our studies demonstrate that phospholipid bilayers catalyze α-syn aggregation at conditions where no aggregates are assembled in bulk solution. The aggregation process was directly observed using time-lapse AFM, showing the number and size of aggregates increasing proportionally with incubation time. The efficient assembly of aggregates on phospholipid bilayers is in line with other studies (Galvagnion et al, 2016; Galvagnion et al, 2015b) in which acceleration of α-syn fibrils formation on phospholipid vesicles was reported. There are a number of important features of this self-assembly process catalyzed by the membrane bilayers.

First, the aggregation efficiency depends on the phospholipid composition, with general aggregation propensity on surfaces following this order: POPS > POPC/POPS > POPC (Table 1). POPS is an anionic phospholipid, suggesting that electrostatic interaction between the negatively charged surface and positively charged segment of α-syn containing lysine residues contributes to the catalytic effect of the POPS surface, in line with (Pfefferkorn et al, 2012). This interpretation is supported by the computational results. Simulations revealed that lysine residues are critically involved in the initial interaction with the membrane surface. Moreover, bilayer composition also contributes to the α-syn structure, which then affects the aggregation propensity of the protein. This is evident in the simulations with membrane-bound and free a-syn molecules; in particular for POPS, the α-syn protein is extended from the bilayer and acts as an attachment point for free proteins to assemble the dimer (Figure 7b and Movie EV4). This extended arrangement is in line with recent structural data (Fusco et al, 2014), according to which, three regions of membrane-bound α-syn exhibit distinct structural and dynamic properties. Thus, that α-syn has differential binding modes on different lipid bilayers, which may alter the overall protein structure and contribute to a change in aggregation propensity of the protein.

Second, it is widely accepted that interaction of amyloid proteins, including α-syn, with lipid bilayers is accompanied with the change of the bilayer structure and even disruption of the bilayer (Jo et al, 2000; Yip et al, 2002; Yip & McLaurin, 2001). The formation of channel-like features assembled by amyloid proteins oligomers is reported in (Quist et al, 2005; Stockl et al, 2013). We have not observed such changes in the bilayer structure in the present study. In Figure 4 aggregate highlighted with blue dissociate and does not appear in frame B, however no damage to the bilayer surface is seen in the images prior and after the aggregate dissociation. This finding suggests that in our experiments the interaction of α-syn takes place with the bilayer groups located on the surface or in the proximity of the surface without insertion of the protein into the bilayer. Computer simulations support these observations, showing that α-syn interaction with POPC and POPS bilayers occur through the lipid head groups and in the interfacial region of the head groups. Explanation can be found in the concentration of amyloid protein used. For example, the α-syn pores in (Quist et al, 2005) were assembled with α-syn concentration three orders of magnitude higher than in our experiments. Another explanation for the discrepancy could be the presence of defects on the bilayer. α-Syn aggregates are reported to sense packing defects, and induce lateral expansion of lipid molecules that progress further to bilayer remodeling by insertion of α-syn into the headgroup region (Ouberai et al, 2013). In our study, we developed a procedure by which the bilayers remain defect-free during the entire time-lapse experiment (Appendix Figure S1). This model is in line with the data of Chaudhary and coworkers (Chaudhary et al, 2016), in which homogeneous bilayers remain intact despite the formation of α-syn oligomers.

Third, the self-assembly of aggregates on the membrane bilayers is a dynamic process. In addition to gradual growing of the aggregate, some of them can dissociate from the surface to the bulk solution (Figure 4). This process leads to the accumulation of aggregates in the solution and direct measurements (Figure 5) support this phenomenon. Our combined experimental and computer modeling approaches demonstrate that the on-surface aggregation is a dynamic process, so the assembled aggregate can dissociate from the surface to the bulk solution. As a result, the dissociated aggregates can play roles of seeds for aggregation in the bulk solution or act as neurotoxic agents. Both processes lead to neurodegeneration. Importantly, in the vast majority of cases, we found that aggregates formed on the surface are oligomers, which are considered to be the most neurotoxic amyloid aggregates.

Based on these studies, we propose the model of amyloid aggregate assembly catalyzed by cellular membranes schematically shown in Figure 8. Interaction of the protein with the membrane changes the protein conformation (panel b), facilitating the interaction with other proteins and assembly of the oligomer (scheme c). The process repeats as more proteins appear leading to the assembly of larger oligomers (scheme d). The assembled oligomer can dissociate from the surface to the intracellular space starting the neurodegeneration effect (scheme e).

**Figure 8.**
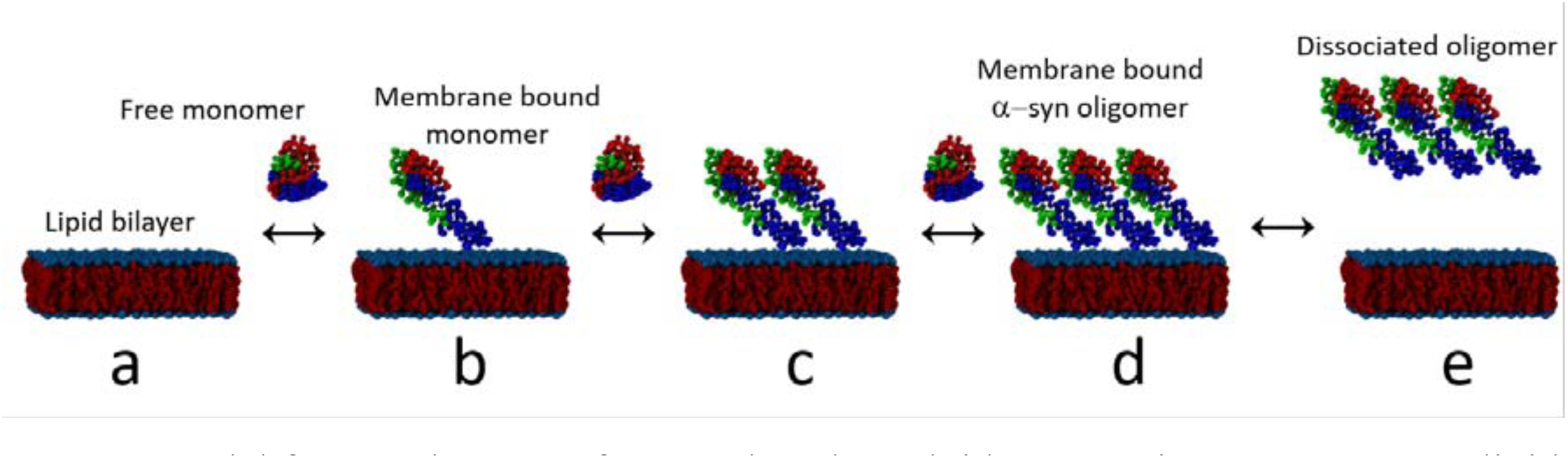
Model for membrane surface catalyzed amyloid aggregation process. **(a)** A lipid bilayer with free α-syn monomers far from the membrane surface. **(b)** Interaction with membrane induces conformation change in the α-syn monomer. **(c)-(d)** The membrane- bound monomer acts as an anchor and attracts free monomers, leading to the formation of oligomers. This process can repeat multiple times, for each repeat the oligomer grows. **(e)** Oligomer dissociates from the membrane to the bulk solution.

One of critical properties of the on-surface aggregation process is that the aggregates form at concentrations as low as the nanomolar range, which corresponds to the typical physiological concentrations of endogenic proteins such as α-syn (Wang et al, 2012). Spontaneous assembly of aggregates in the bulk solution require concentrations several orders of magnitude higher (Bousset et al, 2013), and the amyloid cascade hypothesis considers accumulation of amyloid proteins, which is one of the problem of this model of PD. The problem of the high concentration is alleviated if the assembly occurs on the membrane bilayers. The second important feature of the on-surface aggregation is that the composition of the bilayer contributes to the surface aggregation propensity – namely, a higher anionic lipid content favors α-syn-membrane interactions and lipid-induced α-syn aggregation. Previously reported findings suggest that the levels of anionic lipids in the brain increases with aging (Giusto et al, 2002) and that the ratio of acidic to zwitterionic phospholipids increases in PD brain (Riekkinen et al, 1975). Based on these data and our observations, we hypothesize that amyloidogenic aggregates of α-syn assemble on cellular membrane and the membrane composition is the factor that controls the aggregation process (Ysselstein et al, 2017; Ysselstein et al, 2015). For membranes with normal composition, assembled aggregates are unstable and dissociate as observed in computational modeling of POPC bilayers. A change in the membrane composition, such as switch from POPC rich to POPS rich, leads to a dramatic increase of stability of the dimers, facilitating the assembly of higher order oligomers. Therefore, we posit that the composition of cellular membranes is the factor that defines the healthy state of neurons. Changes in membrane composition leading to an increase in affinity of α-syn for the cell surfaces and favors the formation of stable oligomers and thereby triggers development of the disease.

The proposed model is a significant departure from the current amyloid aggregation model and has two important features. First, it does not require the increase of α-syn synthesis to the level allowing for the spontaneous assembly in aggregates (a few orders of magnitude). Lowering of the protein level is a logical consequence of the drug development efforts in the framework of traditional amyloid aggregation model that did not succeed (Brundin et al, 2017; Busche et al, 2015). Second, α-syn is the protein actively involved in important physiological processes such as the signal transduction in the neuron synapse (Venda et al, 2010). Lowering its level can impair these important processes. In the framework of our model, the protein concentration is not a critical parameter. The property of membrane, such as its ability to facilitate the aggregate assembly mediated by the membrane composition is the factor that defines the disease state, suggesting that preventions and treatments should be focused on the control the membrane composition that can be achieved via controlling the lipid metabolism.

Although the data presented in this paper are obtained for α-syn, the membrane aggregation model can be extended to the amloidogenic proteins and hence to other diseases. The support comes from our recent paper (Banerjee et al, 2017) in which aggregation of α-syn along with the full-size amyloid beta protein (Aβ) on mica surfaces were performed. For both proteins, interaction with the surface dramatically facilitated the aggregation process. Our preliminary data on aggregation of Aβ42 protein revealed a similar catalytic property for aggregation on both POPC and POPS bilayers and hence support our membrane catalyzing model for amyloid aggregation as the molecular mechanism of development of neurodegenerative diseases mediated by protein aggregation.

## Materials and methods

### Materials

1-Palmitoyl-2-oleoyl-sn-glycero-3-phosphocholine (POPC) and 1-palmitoyl-2-oleoyl-sn-glycero-3-phospho-L-serine (POPS) were purchased from Avanti Polar Lipids, Inc, Alabama, US; Chloroform ((> 99.5%, Sigma–Aldrich Inc.); dry bath incubator (Fisher Scientific); A 10 mM pH 7.4 sodium phosphate buffer (PBS, NaH_2_PO_4_•H_2_O: Na_2_HPO_4_ = 1:3.4 without additional salt) was prepared and filtered through a disposable Millex-GP syringe filter unit (0.22 μm) before use. Deionized (DI) water (18.2 MΩ, 0.22 μm pore size filtered, APS Water Services Corp., Van Nuys, CA) was used for all experiments. Muscovite mica (Asheville Schoonmaker Mica Co., Newport News, VA). 1-(3-aminopropyl)silatrane was synthesized as previously described (Rauscher et al); ImmEdge hydrophobic barrier pen (Vector Laboratories, Inc. Burlingame, CA); Aron alpha industrial glue (Toagosei America, West Jefferson, Ohio); S/P Brand Bev-L-Edge micro glass slides (Allegiance Healthcare Corporation, McGaw Park, IL).

### Preparation of α-syn solution

Wild-type A140C α-syn in which the C-terminal alanine was replaced with a cysteine was prepared as described previously (Krasnoslobodtsev et al, 2012). α-Syn solutions were freshly prepared by dissolving 0.4 to 0.8 mg of the lyophilized powder in 200 µL water (the pH was adjusted to 11 with NaOH), in the presence of 1 µL of 1 M dithiothreitol (DTT) to break disulfide bonds, followed by the addition of 300 µL of 10 mM sodium phosphate buffer (pH 7.4). The solution was filtered through an Amicon filter with a molecular weight cutoff of 3 kDa at 14,000 rpm for 15 min. The filtration was repeated 3 times to completely remove free DTT. The concentration of the stock solutions was determined by spectrophotometry (Nanodrop^®^ ND-1000, DE) using the molar extinction coefficients 1280 cm^−1^⋅M^−1^ and 120 cm^−1^⋅M^−1^ for tyrosine and cysteine at 280 nm, respectively. In general, freshly prepared stock solutions were used for all the experiments.

### Preparation of APS-mica

Freshly cleaved mica strips (5.0 × 1.5 cm, L×W.) were immersed in plastic cuvettes containing 167 μM APS solution for 30 min (Krasnoslobodtsev et al, 2012; Lv et al, 2015), followed by rinsing with deionized water and drying in argon flow. The APS-mica was stored in a vacuum chamber for use over the following few weeks (Shlyakhtenko et al, 2013). The APS-mica strips were cut into ∼1.5 ×1.5 cm pieces and glued to a glass slide for sample preparation.

### Preparation of SLBs

We followed a published protocol with minor modifications (Shlyakhtenko et al, 2013). Lipid powder (25 mg) was first dissolved in 1 mL chloroform to make a 25 mg/mL stock solution. The stock solution was aliquoted and stored at −20 °C. The aliquoted solution (20 μL) was thawed and brought to room temperature before it was blow dried with a gentle argon flow and vacuum desiccated overnight. Next, a 1 mL solution of 10 mM sodium phosphate buffer (pH 7.4) was injected into the glass container to make a 0.5 mg/mL solution, unless a different concentration is stated. The solution was sonicated (Branson 1210, Branson Ultrasonics, Danbury, CT) until mostly clear to obtain small unilamellar vesicles. A mica piece mounted on a glass slide was then prepared for experiments by drawing around the perimeter of the mica with a hydrophobic pen to prevent overflow. 60 to 80 μL of the lipid solution was deposited onto the freshly cleaved mica surface. The incubation of lipid solution was done at 60 °C while buffer was supplied periodically. After 1 h, excess lipids were washed off by extensive gentle exchange of the lipid solution with buffer. The resulting SLB was kept in buffer and imaged.

### In situ time-lapse AFM imaging in liquid

AFM imaging was conducted on an MFP-3D (Asylum Research, Santa Barbara, CA). Tapping mode was used. A MSNL cantilever (Bruker, Santa Barbara, CA) with nominal spring constant of 0.1 N/m was used for imaging in liquid. The resonance frequency varied between 7 kHz to 9 kHz. The scan size was 5 μm × 5 μm, and the scan rate was 1 Hz. Buffer was injected periodically to keep the sample at constant volume. For in situ time-lapse AFM experiments, the images were acquired at different time points. Between images, the AFM tip was placed on idle (electronically retracted, approximately 4 μm above the scan area) to ensure that it exerted minimum influence on the sample.

### Data analysis

All AFM images were flattened and then processed using Femtoscan software (Advanced Technologies Center, Moscow, Russia). The features on bilayer surfaces were visually inspected and analyzed using “Grain analysis” in the software. Volumes of aggregates were arranged in histograms and fit with a Gaussian distribution using Origin Pro software (OriginLab, Northampton, MA), yielding Mean ± SD values. Scatter plots of number/volume against incubation times were drawn with the Origin software.

### Molecular dynamics simulations

Lipid bilayers of POPC and POPS were prepared using the *insane.py* script (available at http://md.chem.rug.nl) using the MARTINI2.2refP (de Jong et al, 2013) force field together with the polarizable water (Yesylevskyy et al, 2010) model. The initial bilayer was constructed using, in total, 512 lipids placed randomly in a bilayer structure with 40 water molecules per lipid and 150 mM NaCl. After energy minimization using the steepest decent algorithm, the bilayers were simulated as an NPT ensemble for 500 ns using a 20 fs integration time step. The simulation employed periodic boundary conditions (PBC) with a semi-isotropic pressure coupling using the Parrinello-Rahman barostat at 1 bar with a 12 ps coupling constant. The temperature was kept at 300 K using the velocity rescaling algorithm. Electrostatic interactions were calculated using the particle mesh Ewald algorithm, with a real space cut-off of 1.1 nm. All simulations were performed using the 2016 version of the GROMACS suite of programs (Abraham et al, 2015). Only the final frame of each bilayer simulation was used for further simulations.

Micelle-bound α-syn (PDB ID: 1XQ8) was used as the initial protein structure. A coarse-grained structure was generated using the *martinize.py* script and the PDB structure. The coarse-grained α-syn structure was then placed at a COM distance of 6 nm from the bilayer core in a parallel orientation (along the long protein axis) to the bilayer. The system was then solvated in a box of 13×13×18 nm^3^ water and 150 mM NaCl. The simulation procedure was the same as previously described for bilayers alone, with the exception that the simulation duration was 4 μs for each protein-bilayer system.

Simulations with membrane-bound and additional free α-syn were conducted using the last frame of the 4 μs simulation and adding another α-syn at a COM distance of 6 nm from the membrane-bound α-syn. Orientation of the free α-syn was parallel to the bilayer using the same protein conformation as the initial α-syn-bilayer simulation. Simulations for both POPC and POPS were carried out for 2 μs each using the previously described parameters.

## Acknowledgements

The work at University of Nebraska Medical Center was supported by grants from the National Institutes of Health (NIH) to Y.L.L. (R01 GM096039, R01GM118006 and R21 NS101504). J.C.R. was supported by Branfman Family Foundation. M.H. was partially supported by the UNMC Graduate Fellowship. The computational modeling was partially performed using resources at the Holland Computing Center of the University of Nebraska, which receives support from the Nebraska Research Initiative.

## Author contribution

Y.L.L., M.H. and L.V. designed the project. L.V., K.Z., S.B. performed AFM experiments; M.H. performed the molecular dynamics simulations; J.C.R. provided α-synuclein protein samples. All authors wrote and edited the manuscript.

## Conflict of interest

Authors declare no conflict of interest

## The paper explained

### Problem

Numerous studies have shown that amyloidogenic proteins, including α-syn, are capable of spontaneous assembly into aggregates, and eventually form fibrillar structures found in amyloid or amyloid-like deposits. However, a critical obstacle exists for translating the current knowledge of amyloid proteins aggregation *in vitro* to the aggregation process *in vivo*: the concentration for the spontaneous aggregation of amyloid proteins *in vitro* is in the micromolar range, while physiological concentrations of amyloid proteins are in the low nanomolar range. This nearly thousand-fold difference in concentration suggests the potential role for other cellular factors in promoting aggregation at physiological amyloid proteins concentrations. We hypothesize that self-assembly of the disease-prone amyloid proteins aggregates is driven by the interaction of amyloid proteins with the cellular membrane.

### Results

Consistent with this possibility, we have discovered a novel aggregation pathway in which spontaneous assembly of α-syn protein at the physiological concentration range occurs at the surface rather than in the bulk solution (i.e., an *on-surface* aggregation pathway). Our combined experimental and computer modeling approaches led us to the mechanism of the early stages of protein aggregation in which the key step triggering the disease is the interaction of α-syn monomers with cellular membranes, which catalyzes the aggregation process.

### Impact

Our finding leads to the hypothesis that interaction of amyloid proteins with cellular membrane is the mechanism by which amyloid aggregation can initiate *in vivo* at the physiological concentration range, and the change in membrane properties leading to the increase in affinity of amyloid proteins to the membrane surface triggers of the amyloid aggregation, defining the disease state. Our model is a significant departure from the current model in which amyloid aggregation is linked to elevated synthesis of amyloid proteins. This is a paradigm shift for the development of efficient treatments and diagnostics for protein aggregation in neurodegenerative diseases.

---

**Figure EV1:**
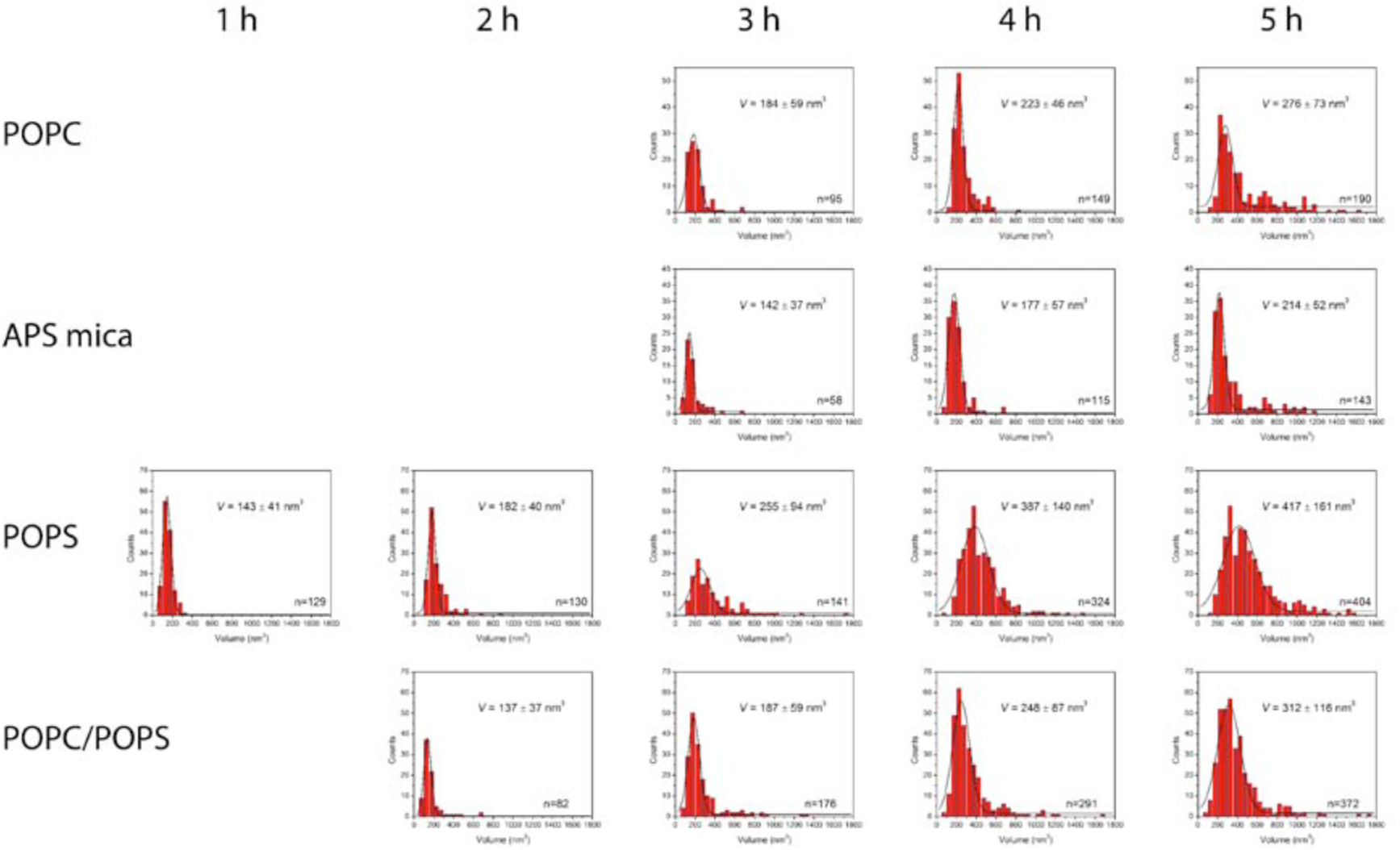
Volume distributions of α-syn aggregates on different surfaces at different time points. Each row represents data from the same surface, while each column is a different incubation time-point. The black curves are Gaussian fits. The most probable volumes are shown as mean ± SD. The number of aggregates is provided in the bottom-right corner of each graph.

**Movie EV1:**
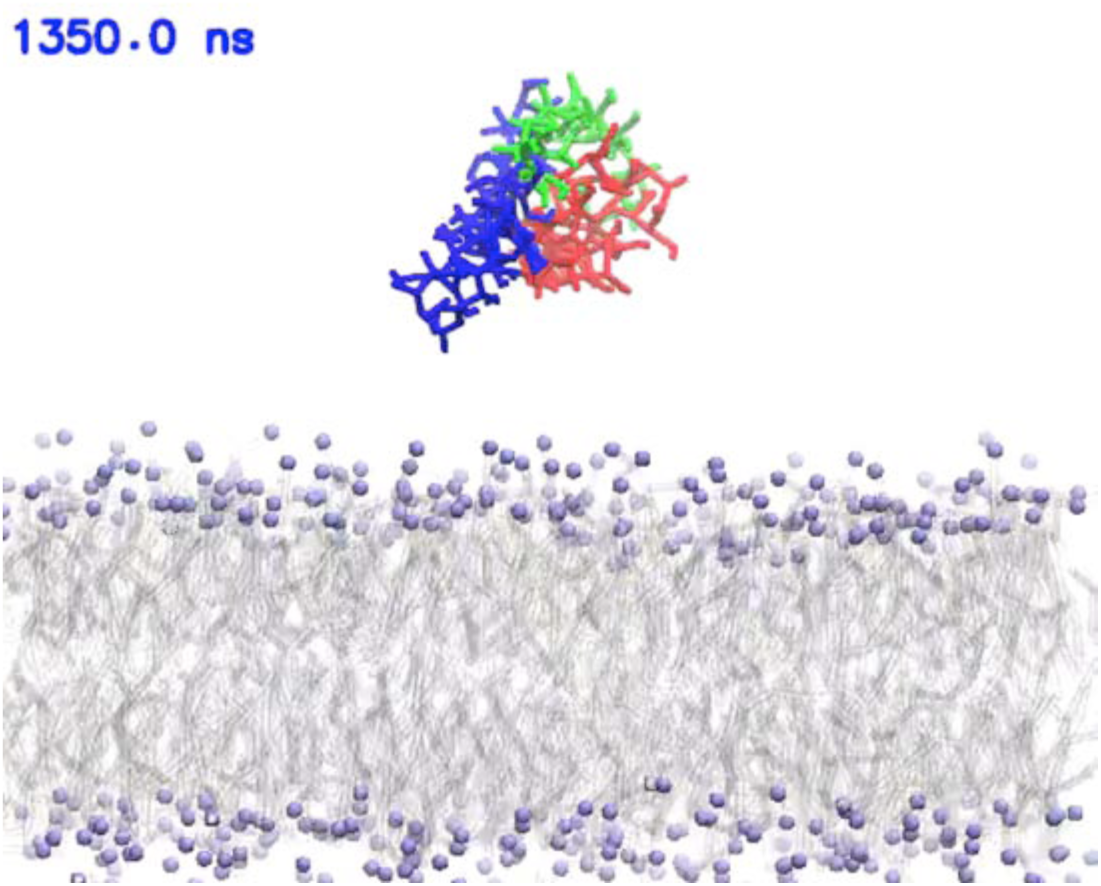
MD simulation of α-syn interaction with a POPC lipid bilayer. Stable binding of the α-syn protein to the bilayer is observed. The α-syn N-terminal segment is colored blue, NAC region is in green, and the C-terminal segment is in red. N- and C-terminal residues are highlighted with a sphere. Lipid tails are in grey, while the POPC head groups are in purple.

**Movie EV2:**
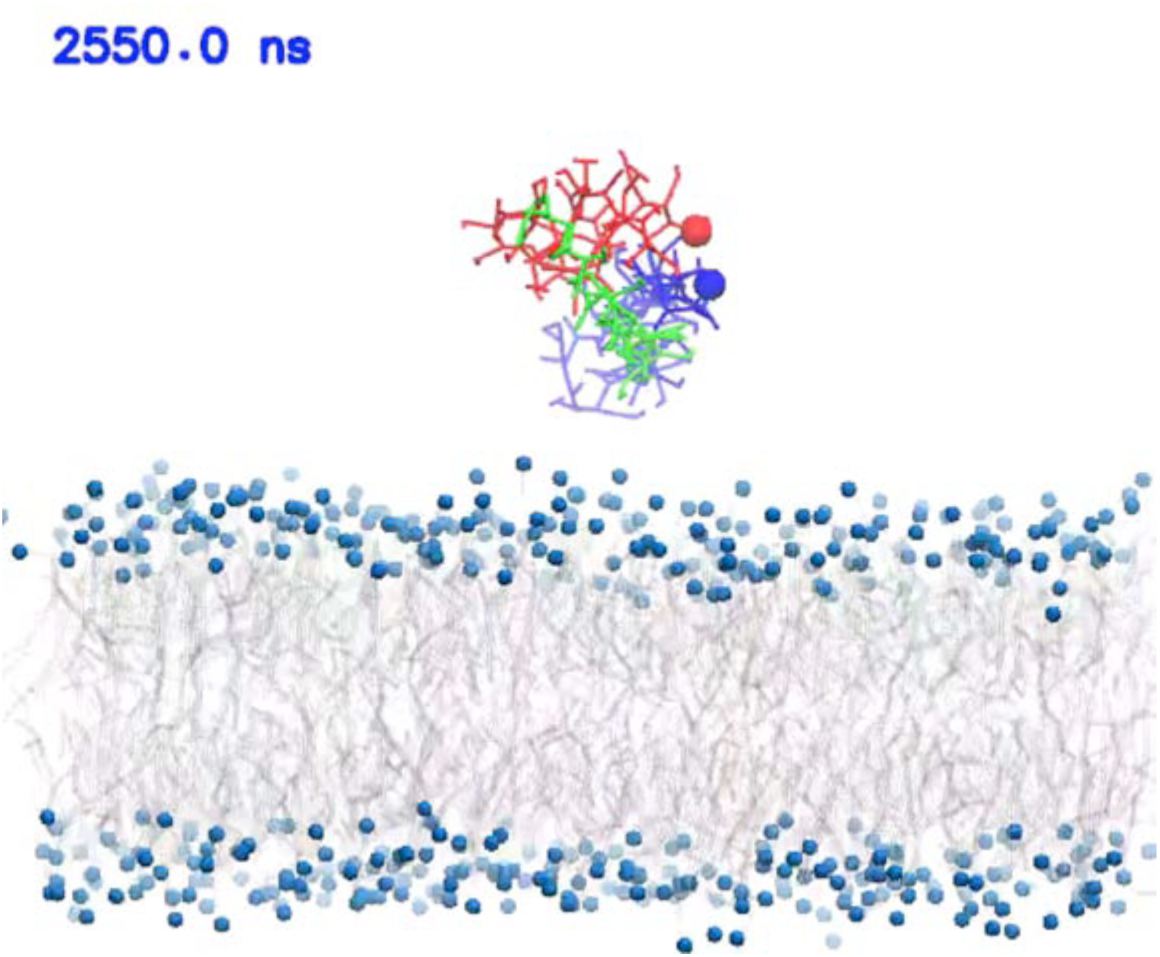
MD simulation of α-syn interaction with a POPS lipid bilayer. The stable binding event of α-syn to the bilayer is shown. α-syn N-terminal segment is colored blue, NAC region is in green, and the C-terminal segment is in red. N- and C-terminal residues are highlighted with a sphere. Lipid tails are in grey, while the POPS head groups are in blue.

**Movie EV3:**
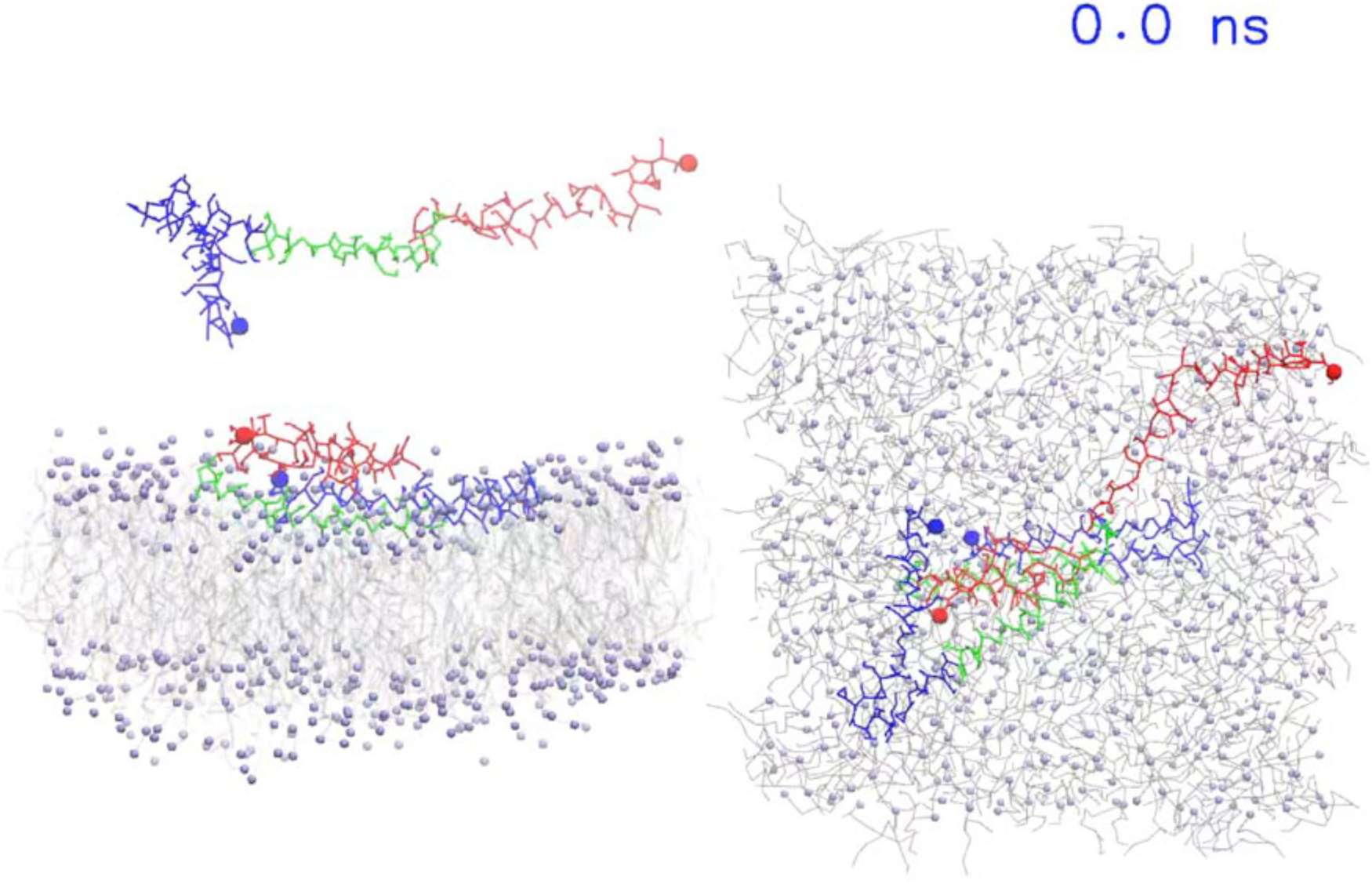
MD simulation of a free α-syn interacting with a membrane bound α-syn on a POPC bilayer. The binding of α-syn to the bilayer is shown. α-syn N-terminal segment is colored blue, NAC region is in green, and the C-terminal segment is in red. N- and C-terminal residues are highlighted with a sphere. Lipid tails are in grey, while the POPS head groups are in blue.

**Movie EV4:**
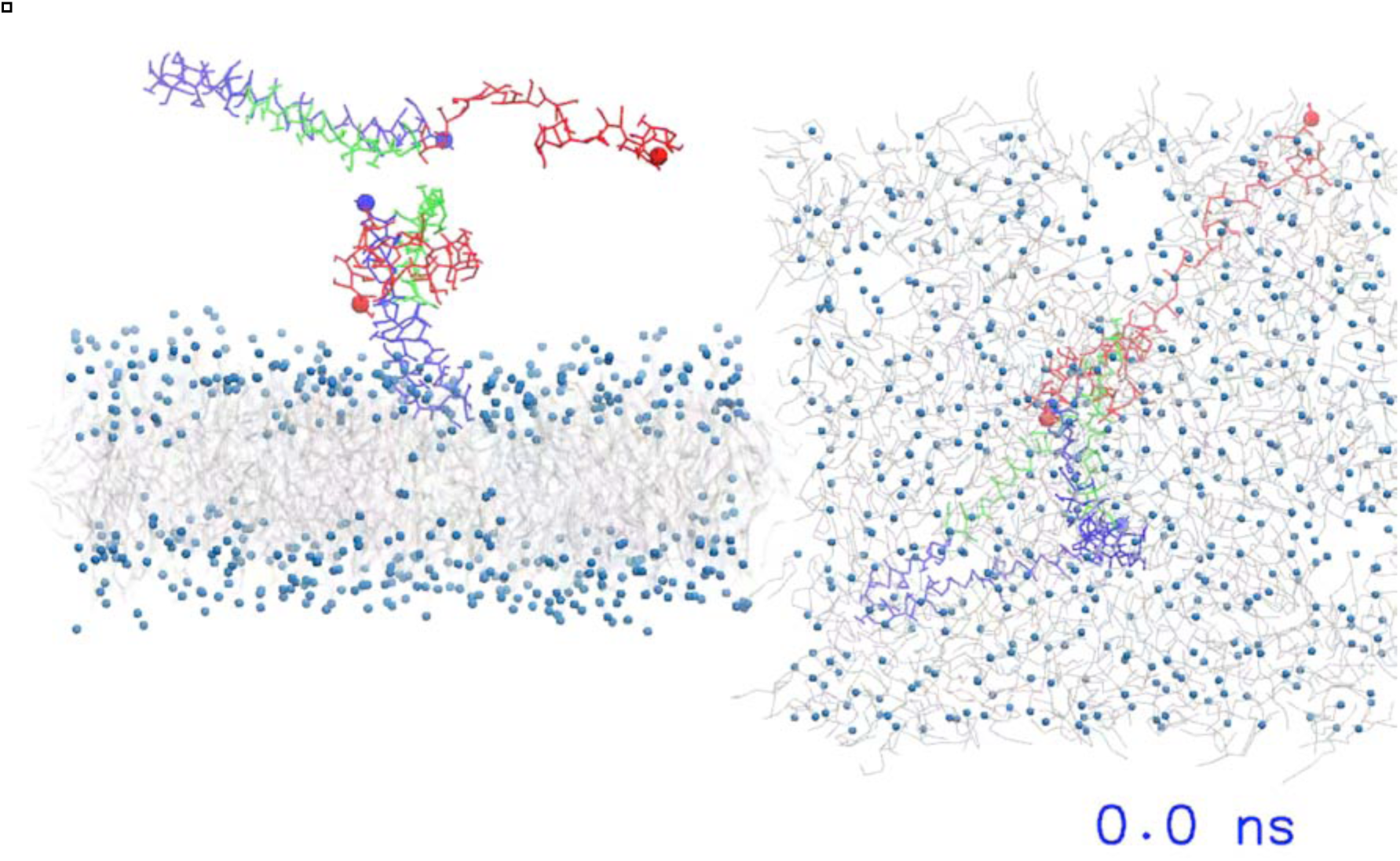
MD simulation of a free α-syn interacting with a membrane bound α-syn on a POPS bilayer. The binding of α-syn to the membrane-bound α-syn is shown. α-syn N-terminal segment is colored blue, NAC region is in green, and the C-terminal segment is in red. N- and C-terminal residues are highlighted with a sphere. Lipid tails are in grey, while the POPS head groups are in blue.

